# Targeting the Autonomic Nervous System Balance in Patients with Chronic Low Back Pain using Transcranial Alternating Current Stimulation: A Randomized, Crossover, Double-Blind, Placebo-Controlled Pilot Study

**DOI:** 10.1101/668541

**Authors:** Julianna H. Prim, Sangtae Ahn, Maria I. Davila, Morgan L. Alexander, Karen L. McCulloch, Flavio Fröhlich

## Abstract

**Background:** Chronic low back pain (CLBP) is characterized by an alteration in pain processing by the central nervous system that may affect autonomic nervous system (ANS) balance. Heart rate variability (HRV) reflects the balance of parasympathetic and sympathetic ANS activation. In particular, respiratory sinus arrhythmia (RSA) solely reflects parasympathetic input and is reduced in CLBP patients. Yet, it remains unknown if non-invasive brain stimulation can alter ANS balance in CLBP patients.

**Objective:** To evaluate if non-invasive brain stimulation modulates the ANS, we analyzed HRV metrics collected in a previously published study of transcranial alternating current stimulation (tACS) for the modulation of CLBP through enhancing alpha oscillations. We hypothesized that tACS would increase RSA.

**Methods:** A randomized, crossover, double-blind, sham-controlled pilot study was conducted to investigate the effects of 10Hz-tACS on metrics of ANS balance calculated from electrocardiogram (ECG). ECG data were collected for 2 minutes before and after 40 minutes of 10Hz-tACS or sham stimulation.

**Results:** There were no significant changes in RSA or other frequency-domain HRV components from 10Hz-tACS. However, exploratory time-domain HRV analysis revealed a significant increase in the standard deviation of normal RR intervals (SDNN) for 10Hz-tACS relative to sham.

**Conclusion(s):** Although tACS did not significantly increase RSA, we found in an exploratory analysis that tACS modulated an integrated HRV measure of both ANS branches. These findings support the further study of how the ANS and alpha oscillations interact and are modulated by tACS.

## Introduction

Chronic pain is a severe, disabling condition that affects 25-30% of the population in the United States (1) and treatment options are limited (2). While opioid therapy has shown short-term efficacy in decreasing pain, few studies have investigated its long-term effectiveness (3) and systematic reviews identify multiple severe risks of long-term use, including: misuse, abuse/dependence, overdose, and death (3,4). Chronic low back pain (CLBP) is the second most prevalent cause of disability in adults in the US (5). The poor rates of recovery (58% at 1 month) and high rates of recurrence (73% in 12 months) contribute to high social and economic costs (6).

CLBP often persists without clear peripheral pathology (peripheral injury may trigger but does not sustain CLBP) and the mechanism of pain development is not fully understood (7). In CLBP, the relationship between nociception and pain is often weak or lost indicating abnormal integration (8), which points to an alteration in pain processing by the central nervous system (9,10). CLBP stems from dynamic interactions between sensory and contextual (i.e., cognitive, emotional, and motivational) processes in the brain that are mediated by feed-forward and feedback processes (8). Recent neurobiological investigations support the crucial role of the brain within chronic pain development by showing substantial structural, physiological, and metabolic changes (8) including autonomic nervous system (ANS) balance (11). The ANS controls a range of vital involuntary physiological functions, such as regulating blood pressure, temperature, and heart rate at rest and in response to stressors (12). Regulatory ANS function can be quantitatively assessed by the analysis of the heart rate variability (HRV), which is the variation in time between successive heartbeats. HRV is composed of input from the excitatory sympathetic and inhibitory parasympathetic nervous system as well as baroreceptors and vagal tone. When HRV is deconstructed through signal processing, it is possible to quantify rhythmic components that reflect specific pathways of the ANS neural regulation. The most salient components are a respiratory oscillation known as respiratory sinus arrhythmia (RSA)(13) and low-frequency (LF) components assumed to be related to blood pressure regulation via the baroreceptors and peripheral vasomotor activity (14,15).

Pain signal regulation is a normal part of the defensive response mediated by the nervous system. The body reacts to illness by activating and sensitizing afferent nociceptive neurons (16). In the case of chronic pain, this process may trigger hyperarousal of the sympathetic nervous system (17). Based on the Polyvagal theory (18,19), an evolutionary neurophysiological model of the autonomic response to safety and threat, chronic maintenance of threat responses can lead to a compromised functional state (20). These chronic systemic functional problems are reflected in the regulation of the heart by the most rapidly responsive component of the nervous system, the ventral vagal complex, as measured by RSA (19). Previous studies have targeted the reduced RSA using biofeedback interventions, and HRV components have been used to measure the efficacy of chronic pain therapies (21–24). Yet, little is known if targeting network pathologies by non-invasive brain stimulation can influence ANS balance in patients with CLBP.

We performed a randomized, crossover, double-blind, sham-controlled clinical trial to investigate the effect of transcranial alternating current stimulation (tACS), which is a form of non-invasive brain stimulation that applies weak sine-wave electrical current to the scalp and can modulate oscillatory brain network activity (25–27), in patients with CLPB. In a separate publication, we reported that tACS enhanced impaired alpha oscillations and that pain relief correlated with the stimulation-induced increase in alpha oscillations in patients with CLBP (28). We hypothesized that tACS would increase RSA, and therefore reduced CLBP, based on previous findings that patients with CLBP show pathologically reduced RSA compared to healthy controls (29–34).

## Methods

### Participants

The inclusion criteria were as follows: (1) age between 18 and 65; (2) diagnosis of chronic low back pain by a clinician; (3) pain for at least 6□months; (3) an average daily pain rating ≥4 as measured by a 0-10 numeric rating scale (NRS); (4) no history of neurologic or psychiatric conditions and no current unstable medical conditions; (5) no contraindications to tACS; and (6) no current pregnancy. The study was approved by the Biomedical Institutional Review Board of University of North Carolina at Chapel Hill and registered on clinicaltrials.gov (NCT03243084). Participants were recruited from local pain and physical therapy clinics, as well as the University listserv email and a recruitment website (jointheconquest.org). Participation consisted of two sessions and two follow up emails after a telephone screening determined initial eligibility. Participants also met criteria for low depression (total score <17) and suicide risk as defined by the Hamilton Depression Rating Scale (35) (suicide question score ≤2).

### Study Design

We conducted a randomized, crossover, double-blind, sham-controlled trial. Participants received both 10Hz-tACS and active sham stimulation in a randomized and counter-balanced order with a separation of at least one week between sessions. Each stimulation session was preceded and followed by ECG, clinical assessments, and electroencephalogram (EEG).

### Brain Stimulation

We applied 10Hz-tACS via three silicone-carbon electrodes on the scalp with Ten20 conductive paste (Bio-Medical Instruments, Clinton Township, MI) and the XCISTE 100 stimulator (Pulvinar Neuro, Chapel Hill, NC). The two stimulation electrodes (5×5cm) were placed at F3 and F4 according to the 10-20 international coordinate system (Figure 1). Stimulation montage and modeling of electric field distribution were calculated by the tES LAB 1.0 software (Neurophet Inc., Seoul, South Korea). The return electrode (5×7cm) was placed slightly below Pz. The two stimulation electrodes each delivered an in-phase sinusoidal waveform with 1mA zero-to-peak amplitude. Stimulation ramped up and down for 10 seconds. For active 10Hz-tACS, the stimulation lasted for 40 minutes. Sham stimulation was identical to active, except that stimulation only lasted for one minute. All participants completed the 10Hz-tACS and sham stimulation for 40 minutes on a different day. There was a required gap of at least 7 days between the two sessions to reduce carry-over effects (14.4 days ± 6.5). Five-digit codes were used to ensure that study coordinators were blind to the stimulation condition. During stimulation, all participants were seated comfortably and watched Reefscapes (Undersea Productions, Queensland, Australia), whichdisplays tropical fish in underwater scenes, to minimize the phosphenes induced by stimulation. Participants were asked to stay relaxed, watch the video, and keep their eyes open.

**Figure 1.**
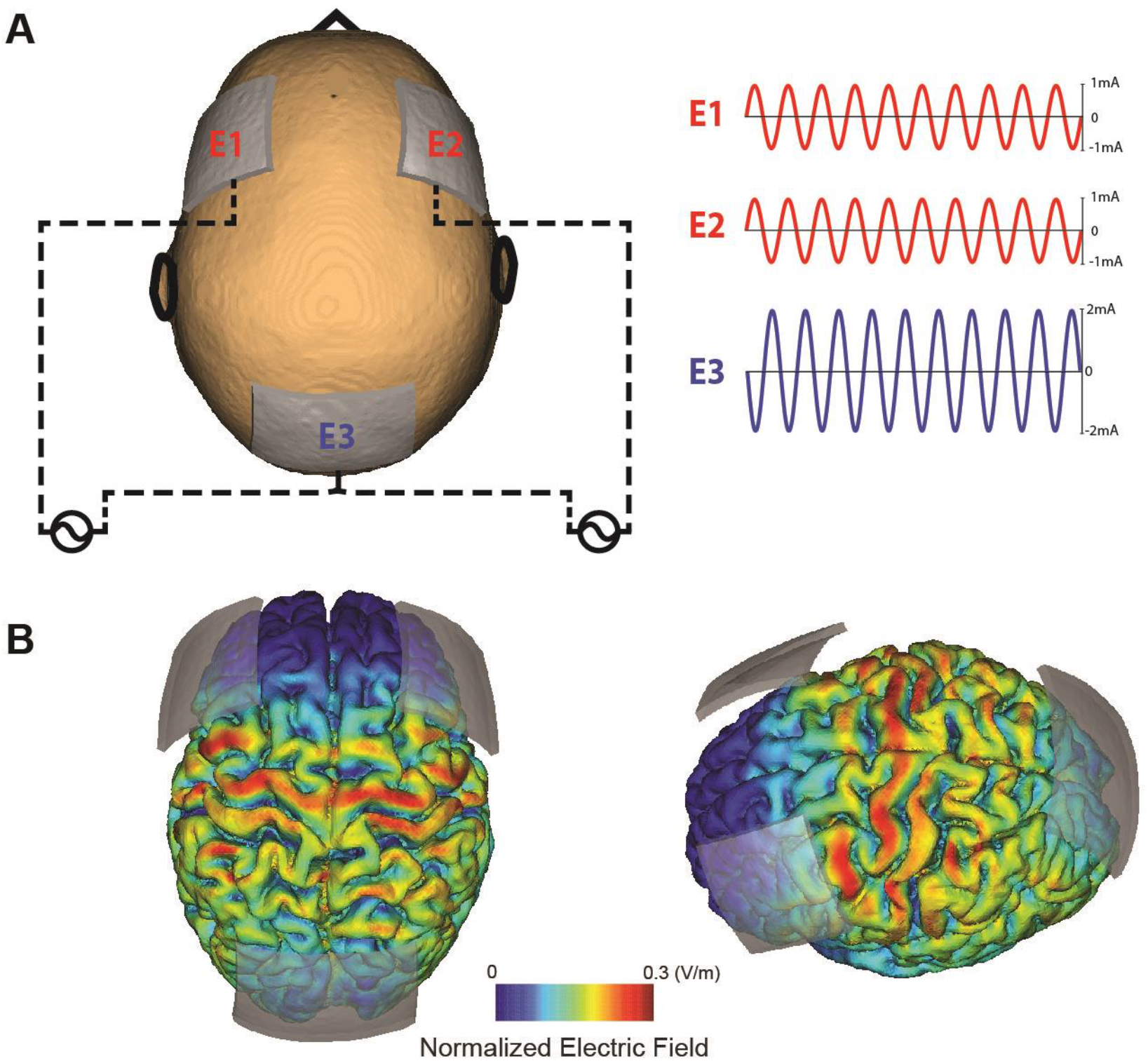
Stimulation montage and electric field distribution. (A) Electrode E1 and E2 deliver 1mA (zero-to-peak) in-phase sine-wave. Electrode E3 is used as a return electrode. (B) Normalized electric field distribution on the cortex (left: top-view, right: left-view).

### Clinical Assessments

Participants completed a battery of baseline surveys including demographics, handedness, State-Trait Anxiety Inventory (STAI) (Trait-version)(36), Behavioral Inhibition and Activation Scales (BIS/BAS)(37,38), and pre-treatment opinion on the likelihood of pain improvement (0-10 numeric rating scale). Other self-report baseline measures included the Pain Catastrophizing Scale efficacy Questionnaire (PSEQ)(43), which assessed pain experience, depression levels, physical activity limitations, and confidence in daily activities respectively. Pain severity and disability were assessed both prior to and after receiving stimulation. The pain scale utilized was an 11 point NRS (0-10) that includes word and facial descriptions and is part of the previously validated Defense and Veterans Pain Rating Scale (DVPRS) (44). The DVPRS was a repeated measure completed at the beginning and end of both sessions and a two-day follow up email. It also includes domain specific questions about pain interference in activity, mood, sleep, and stress in the last 24 hours (answered at session beginning and follow up only). Disability was measured by the Oswestry Disability Index (ODI) (45,46), a back pain specific assessment measuring perception of disability. The ODI was another repeated measure that was completed at the beginning of each session and at follow-up. A pressure pain threshold (PPT) test using the Wagner FDX Algometer (Wagner Instruments FDX-25, Greenwich, Connecticut) was assessed to help quantify and document levels of pain sensitivity via pain tolerance measurement. PPT was assessed before and after stimulation at the right brachioradialis and right sacroiliac joint (47). The participant was instructed to inform the assessor when they first perceived a sensation of pain. The amount of pressure in pounds (lb) that constituted the pain sensation was recorded. This process was repeated three times bilaterally at each site and the average of these measures was used in the data analysis. The test-retest reliability of PPT measurements has been established in previous studies. (48,49).

### Data Collection and Analysis

ECG data were collected with the Physio16 input box (Geodesic EEG System, EGI Inc., Eugene, OR) at a sampling rate of 1kHz. HRV components were extracted from ECG data (14,50) to evaluate changes in neural regulation of the ANS before and after intervention. We placed two disposable snap electrodes below the right collarbone and left inferior costal arch. Inter-beat intervals (IBI), which is the time between consecutive heartbeats expressed in milliseconds, were derived from detected R peaks in ECG data. The R peaks were extracted using the Cardio Peak-Valley Detector (CPVD) (51) and the IBI event series were obtained. The unedited IBI file was visually inspected and edited offline with CardioEdit software (developed in the Porges laboratory and implemented by researchers trained in the Porges laboratory). Editing consisted of integer arithmetic (i.e., dividing intervals between heart beats when detections of R-wave from the ECG were missed or adding intervals when spuriously invalid detections occurred). The resulting normal RR intervals were used in analysis when abnormal beats, like ectopic beats (heartbeats that originate outside the right atrium’s sinoatrial node) have been removed. (52). HRV time and frequency components were calculated with MATLAB and CardioBatch software (Brain-Body Center, University of Illinois at Chicago), respectively. For HRV time components, the average of normal RR intervals (meanRR), the standard deviation of normal RR intervals (SDNN), and the root mean square differences of successive RR intervals (RMSSD) were calculated using custom-built scripts in MATLAB. For HRV frequency components, RSA was calculated using CardioBatch software, which implements the Porges-Bohrer metric (50). This metric is neither moderated by respiration, nor influenced by nonstationarity, and reliably generates stronger effect sizes than other commonly used metrics of RSA (steps are described in depth in (53) and validated in (50). To derive the other HRV frequency components (LF, and LF/HF), the IBI event series was resampled at 2 Hz to generate an equally spaced intervals time series. RSA and LF were calculated based on the Porges-Bohrer method (53,54); RSA uses a third-order, 21 point moving polynomial filter (MPF) applied to the 2 Hz IBI time series to remove low frequency oscillations and slow trend. The residual detrended output of the MPF was filtered with a Kaiser FIR windowed filter with cut-off frequencies that removes variance not related to spontaneous breathing in adults (0.12 to 0.40 Hz). The filtered detrended output was divided into sequential 30-second epochs and the variance within each epoch was transformed by a natural logarithm (ln(ms2)), the mean of these epoch values was used as the estimate of RSA for the specific segment. LF uses a third-order, 51 point moving polynomial filter (MPF) applied to the 2 Hz IBI trend to remove extremely low frequency oscillations and slow trend. The residual detrended output of the MPF was filtered with a Kaiser FIR windowed filter with cut-off frequencies (0.04 to 0.10 Hz). The filtered detrended output was divided in 30 second epochs and the variance within each epoch was transformed with a natural logarithm (ln(ms2)), the mean of the epochs values was used as an estimate of LF for the segment(55). These variables included:1) respiratory sinus arrhythmia (i.e., RSA or high frequency HRV defined by the frequencies of spontaneous breathing (.12-.4 Hz), 2) low frequency HRV (i.e., occurring within the frequencies of spontaneous vasomotor and blood pressure oscillations; .06-.10 Hz), and 3) ratio of LF and HF (LF/HF)

### Statistical Analyses

All statistical analyses were performed using custom-built scripts in R (R Foundation for Statistical Computing, Vienna, Austria). All analyses were run on the difference of data before and after stimulation (post - pre) after taking the natural logarithm (within Porges-Bohrer method). A two-way repeated-measures ANOVA was run with a factor for the stimulation condition (10Hz-tACS vs. sham) and session (first vs. second) for all other HRV components. The session factor was included to control for non-specific effects that the session order might induce.

## Results

### Demographics

The consort diagram is presented in Figure 2 (28). Twenty of the twenty-one participants recruited completed the study. Eighty percent of participants reported CLBP with a duration of greater than two years (Table 1). The most common previous treatment reported included the use of NSAIDs, physical or aquatic therapy, alternative treatments, and low impact exercise such as yoga. All participants reported trying at least two previous treatment options. Full demographics are reported in Table 1.

**Table 1.** Demographic Information. BMI: Body Mass Index, BIS/BAS: Behavioral Inhibition System/ Behavioral Activation System Scale, HamD-17: Hamilton Depression Rating Scale, HRV: Heart Rate Variability, LF/HF: ration of Low Frequency to high frequency, Mean RR: Mean time between RR (all R peaks) intervals, Baseline refers to the pre-stimulation measures of session 1.

| Demographics | Participants (n=20) |
| --- | --- |
| Age | 43.00(13.37) |
| Sex, N(%) |  |
| Male | 8(40%) |
| Female | 12(60%) |
| Race, N(%) |  |
| Caucasian | 18(90%) |
| African American | 2(10%) |
| Ethnicity, N(%) |  |
| Hispanic/Latino | 1(5%) |
| Non-Hispanic/Latino | 19(95%) |
| BMI | 25.94(4.46) |
| Handedness |  |
| Right | 18(90%) |
| Left | 2(10%) |
| Time in Pain (years) | 7.1(6.0) |
| 0-2 yrs, N(%) | 4(20%) |
| 2-5 yrs N(%) | 8(40%) |
| 5+ yrs. N(%) | 8(40%) |
| Previous Treatment, N(%) |  |
| Physical or Aquatic therapy | 13(65%) |
| Opioids | 4(20%) |
| Over the counter medications (e.g., NSAID) | 18(90%) |
| Alternative Treatments (e.g., chiropractor, acupuncture) | 13(65%) |
| Surgery | 3(15) |
| Counseling, Cognitive Behavioral Therapy | 1(5%) |
| Social support (e.g., chronic pain social group) | 1(5%) |
| Low-impact exercise (e.g., yoga, pilates) | 13(65%) |
| Mindfulness Intervention | 3(15%) |
| Other | 5(25%) |
| **Self-Report Assessments (Baseline)** |  |
| Pre-treatment Opinion on likelihood of pain improvement (0-10 NRS, not likely at all to very likely) | 3.75(1.62) |
| Defense and Veterans Pain Rating Scale | 4.4(1.05) |
| Pain Interference with: |  |
| Activity | 4.8(1.67) |
| Sleep | 5.00(2.62) |
| Mood | 4.10(1.68) |
| Stress | 4.35(2.23) |
| Oswestry Disability Index | 28.52(11.51) |
| UCLA Activity score | 5.9(2.29) |
| Low, N(%) | 4(20%) |
| Medium, N(%) | 10(50%) |
| High, N(%) | 6(39%) |
| Pain Catastrophizing Scale (PCS) Total | 15.05(9.78) |
| Rumination | 5.8(4.76) |
| Magnification | 3.2(2.38) |
| Helplessness | 6.05(3.65) |
| Pain Self Efficacy Questionnaire (PSEQ) Total | 37.6(12.14) |
| BIS/BAS Scale |  |
| BAS Drive | 10.3(2.36) |
| BAS Fun Seeking | 11.9(1.97) |
| BAS Reward Responsiveness | 17.3(1.95) |
| BIS | 20.3(4.12) |
| State-Trait Anxiety Inventory (STAI-Trait) | 43.25(10.36) |
| Depression and Anxiety Subscale (DASS-21) | 15.55(13.34) |
| **Clinical Assessments (Baseline)** |  |
| HAMD-17 | 8.9(3.46) |
| Pressure Pain Threshold (PPT) |  |
| Brachioradialis (lb) | 4.8(2.37) |
| Sacroiliac Joint (lb) | 8.99(3.95) |
| **HRV Components (Baseline)** |  |
| Respiratory Sinus Arrhythmia (RSA) or High Frequency (HF) | 5.45(1.41) |
| Low Frequency (LF) | 4.84(1.37) |
| LF/HF | 0.89(0.21) |
| Mean RR | 899.99(149.04) |
| Standard Deviation of RR Intervals (SDNN) | 46.56(27.63) |
| Root Mean Square of Successive Differences (RMSSD) | 41.02(36.58) |

**Figure 2.**
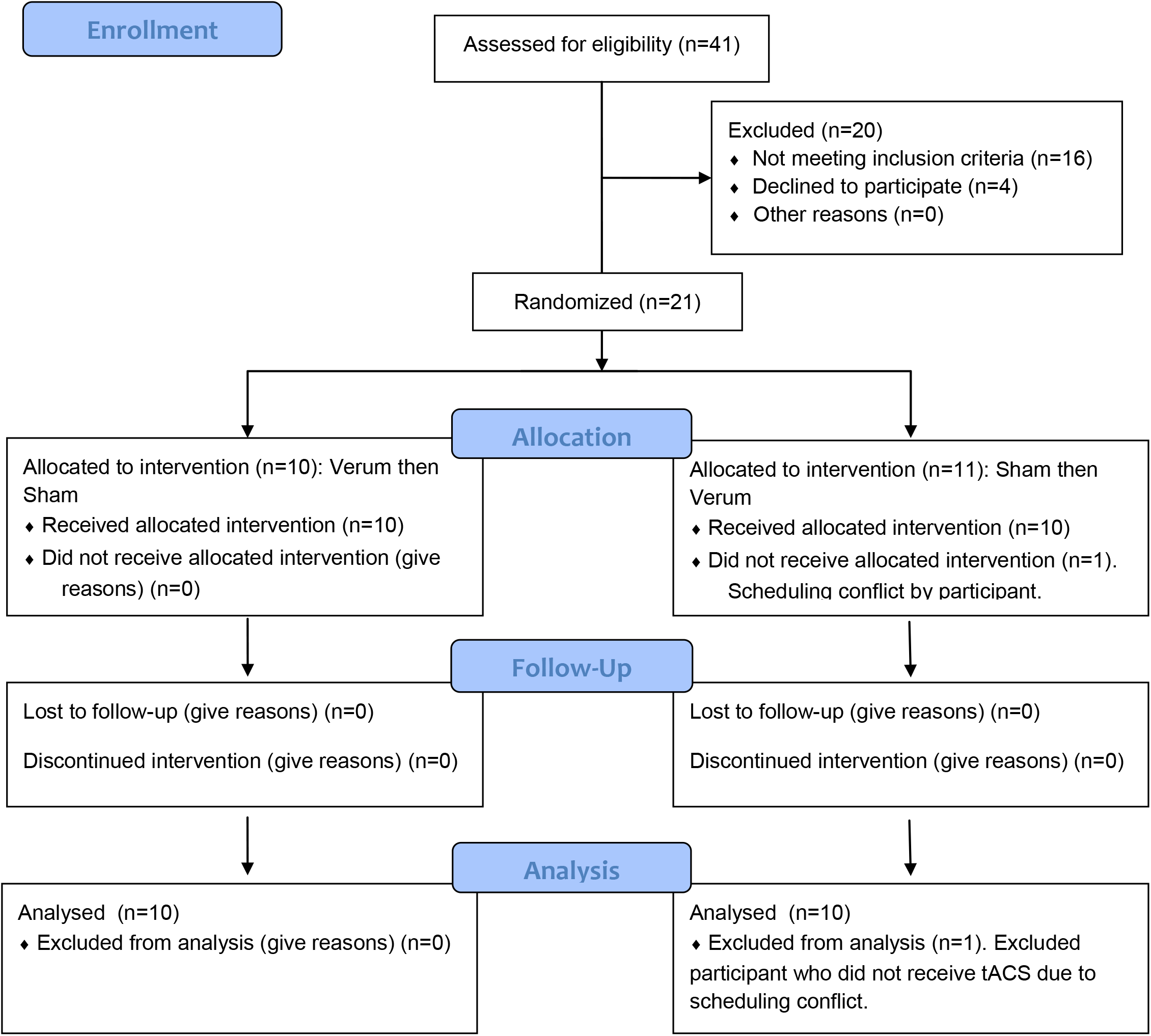
Consort (2010) flow diagram.

### RSA and HRV components

The HRV frequency-domain components followed normal distribution as defined by the Wilks-Sharpiro test (p>.05). RSA was analyzed using a two-way repeated-measured ANOVA of condition (10 Hz-tACS and sham) and session (first visit and second visit). In this analysis the interaction of condition and session was considered to represent a sequence effect if present. For RSA we found no significant main effect for condition (F_1,35_ = 1.01, p =0.32), session (F_1,35_ = 0.33, p = 0.57), or sequence (F_1,35_ = 0.66, p = 0.42) (Figure 3A, left panel). We ran the same two-way repeated-measures ANOVA for LF and LF/HF. We found no significant effects of condition (F_1,35_=1.56, p=0.22), session (F_1,35_=2.91, p=0.10), or sequence (F_1,35_ = 1.44, p = 0.24) for LF nor did we find significant effects of condition (F_1,35_=0.14, p=0.72), session (F_1,35_=2.63, p=0.11), or sequence (F_1,35_ = 0.84, p = 0.37) for LF/HF (Figure 3A, middle and right panels). In the time-domain analyses, we found a trending significant main effect of condition for meanRR (F_1,35_=2.92, p=0.096) but not session (F_1,35_=0.012, p=0.91) or sequence (F_1,35_=0.11 p=0.74; Figure 3B, left panel). For the standard deviation of normal RR intervals (SDNN), we found a significant main effect of condition (F_1,35_=5.34, p=0.027), but not session (F_1,35_=0.00, p=0.999) or sequence (F_1,35_=0.29, p=0.59; Figure 3B, middle panel). Post-hoc paired t-tests yielded a significant SDNN increase in the 10Hz-tACS condition (t_19_=2.07, p=0.05). We found no significant difference in RMSSD for condition (F_1,35_=2.59, p=0.12), session (F_1,35_=0.16, p=0.69), or sequence (F_1,35_=0.02, p=0.88; Figure 3B, right panel). These findings suggest that tACS modulated total HRV (both sympathetic and parasympathetic input) in patients with CLBP. Values for the pain and HRV metrics are presented in Table 2.

**Table 2.** Pre and Post Stimulation Metrics between Treatment Groups for Primary Pain and all HRV Outcome Variables. Each group: N=20 unless otherwise noted. *: n=19, ^+^:n=17.

| Variable | Time point | Treatment | Mean(SD) |
| --- | --- | --- | --- |
| DVPRS (self-report pain) | Pre | 10Hz-tACS | 4.20(1.36) |
|  |  | Sham tACS | 3.95(1.35) |
|  | Post | 10Hz-tACS | 3.10(1.45) |
|  |  | Sham tACS | 3.45(1.77) |
|  | 2 day follow-up | 10Hz-tACS<sup>#</sup> | 3.37(1.26) |
|  |  | Sham tACS<sup>+</sup> | 3.24(1.31) |
| Respiratory Sinus Arrhythmia (RSA) or High Frequency (HF) | Pre | 10Hz-tACS | 5.99(1.35) |
|  |  | Sham tACS | 5.12(1.75) |
|  | Post | 10Hz-tACS | 5.98(1.04) |
|  |  | Sham tACS | 5.32(1.61) |
| Low Frequency (LF) | Pre | 10Hz-tACS | 5.15(1.56) |
|  |  | Sham tACS | 4.86(1.79) |
|  | Post | 10Hz-tACS | 5.21(1.08) |
|  |  | Sham tACS | 5.28(1.81) |
| LF/HF | Pre | 10Hz-tACS | 0.85(0.17) |
|  |  | Sham tACS | 0.95(0.25) |
|  | Post | 10Hz-tACS | 0.87(0.12) |
|  |  | Sham tACS | 1.00(0.25) |
| Mean RR | Pre | 10Hz-tACS | 879.54(175.50) |
|  |  | Sham tACS | 881.91(126.23) |
|  | Post | 10Hz-tACS | 851.57(174.90) |
|  |  | Sham tACS | 911.18(151.64) |
| Standard Deviation of RR Intervals (SDNN) | Pre | 10Hz-tACS | 40.07(23.46) |
|  |  | Sham tACS | 48.23(30.17) |
|  | Post | 10Hz-tACS | 48.14(27.03) |
|  |  | Sham tACS | 48.55(28.38) |
| Root Mean Square of Successive Differences (RMSSD) | Pre | 10Hz-tACS | 34.56(28.29) |
|  |  | Sham tACS | 42.08(36.31) |
|  | Post | 10Hz-tACS | 39.79(34.54) |
|  |  | Sham tACS | 42.87(37.88) |

**Figure 3.**
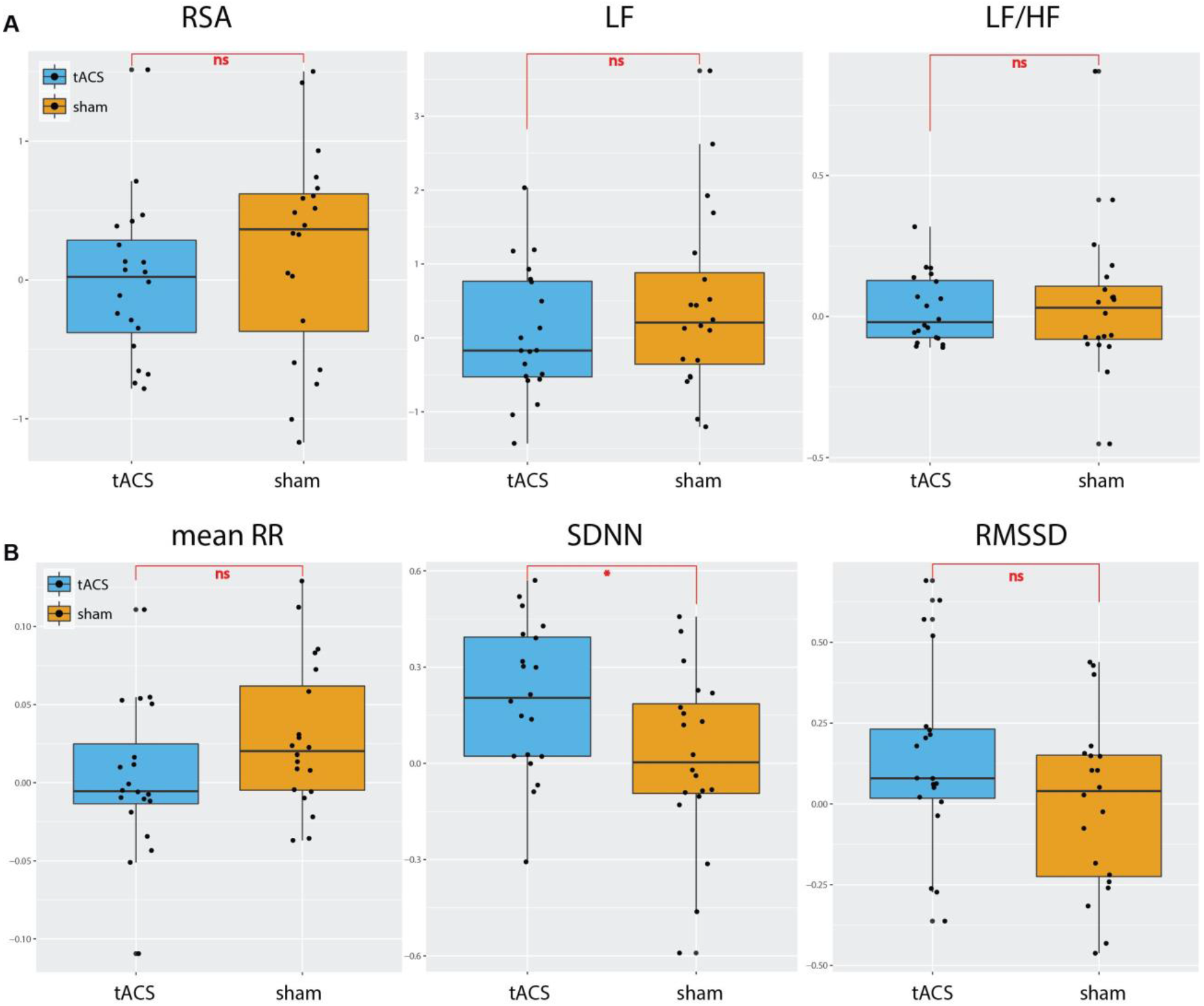
HRV changes [ln(post)-ln(pre)] between tACS and sham conditions for (A) Frequency-domain HRV components; RSA, LF, and LF/HF and for (B) Time-domain HRV components; meanRR, SDNN, and RMSSD. (*:p<0.05, ns: not significant)

### HRV Correlation to Pain

To investigate the presence of a relationship between baseline RSA and pain level, we calculated the Pearson correlation for the pre-stimulation time point (Table 3A). Neither RSA nor SDNN measures correlated with baseline pain level, ODI, or PPT (Table 3A). We also investigated the change in RSA and SDDN with pain, ODI, and PPT change. We found neither RSA nor SDNN correlated with pain, ODI, and PPT change (Table 3B).

**Table 3.** Correlations of HRV Measures (RSA and SDNN) with Pain Measures. a) Correlations at baseline. Baseline refers to the pre-session measures of session 1., b) Correlations for the change (post-pre stimulation) at both sessions. N=20 participants. RSA-Respiratory Sinus Arrhythmia, SDNN- Standard Deviation of NN intervals, DVPRS- Defense and Veterans Pain Rating Scale, PPT- Pressure Pain Threshold, ODI- Oswestry Disability Index, ll All Pearson Correlations, all p-values >.10

| A) | Changes (Post-Pre) | RSA |  | SDNN |  |
| --- | --- | --- | --- | --- | --- |
|  |  | Pearson | p-value | Pearson | p-value |
|  | DVPRS | .080 | .643 | -.247 | .125 |
|  | ODI | .060 | .712 | -.060 | .737 |
|  | Brachioradialis (lb) PPT | .102 | .530 | .224 | .164 |
|  | Sacroiliac Joint PPT | -.058 | .720 | -.230 | .153 |

| B) | Baseline | RSA |  | SDNN |  |
| --- | --- | --- | --- | --- | --- |
|  |  | Pearson | p-value | Pearson | p-value |
|  | DVPRS | -.139 | .557 | .057 | .812 |
|  | ODI | .140 | .559 | -.169 | .477 |
|  | Brachioradialis (lb) PPT | -.363 | .116 | -.002 | .993 |
|  | Sacroiliac Joint PPT | -.132 | .578 | -.127 | .592 |

### Responders

Responders in this study are defined by participants who had a decrease of two points or more on the DVPRS (11 point NRS) after stimulation, which indicates the minimal clinically important difference (MCID) in CLBP based on previously reported findings (56). Twice as many participants reported being a responder (≥2 decrease in pain scale) in the 10Hz-tACS versus the sham condition (8 of responders in 10Hz-tACS vs. 4 of responders in sham). A chi-square test of independence was performed to examine the relation between stimulation condition and being a responder. The relation between these variables did not reach statistical significance, χ^2^ (2, *N* = 2) = 14.14, *p* =0.15, with a high Odd’s Ratio (OR=2.67).

### Blinding and Side Effects

Participants were asked how sure they were of having received stimulation on a visual analog scale (0-100). A t-test was used to analyze confidence in stimulation. There was no significant difference between conditions (t(36.74)=1.38, p=0.18), therefore blinding was considered successful. All participants completed a side-effect questionnaire after both sessions and there were no significant differences in any of the queried potential side-effects between active 10Hz and sham conditions (Table 4).

**Table 4.**
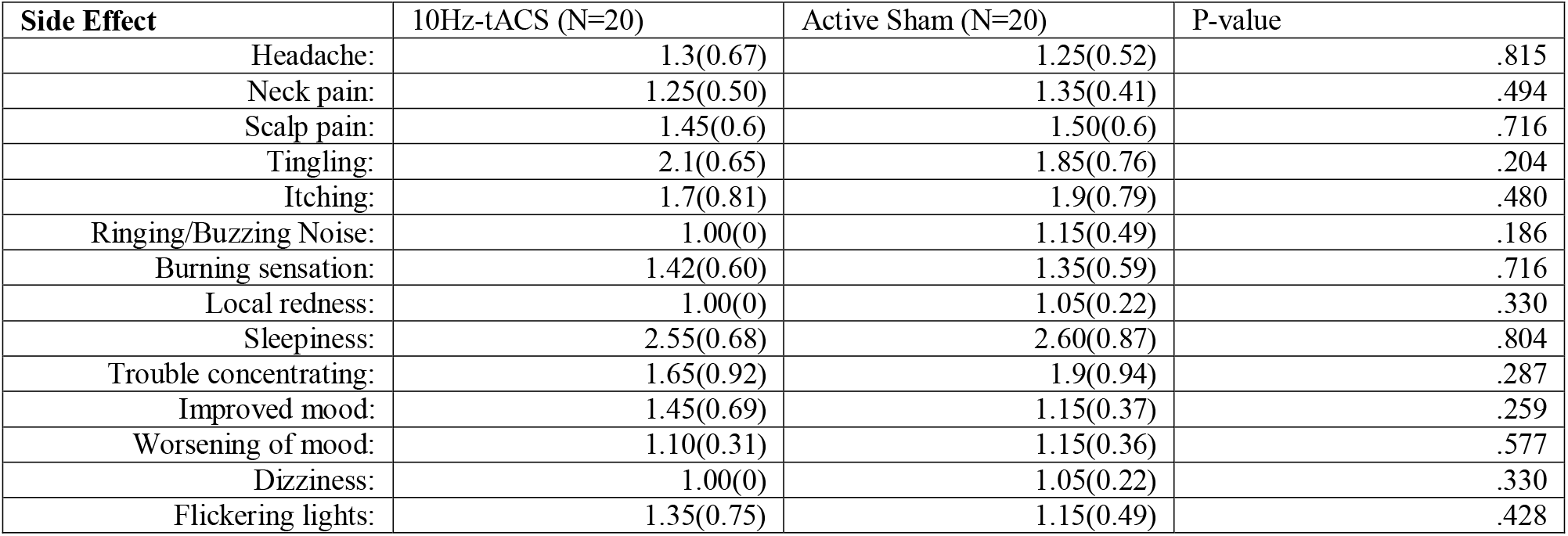
Side Effect Differences Between Conditions. A side effects questionnaire was completed after stimulation at both sessions with 1=absent, 2=mild, 3=moderate, 4=severe. The Mean(SD) are reported for both conditions and paired t-tests were used to test for differences between groups. (*p<.05).

## Discussion

In this study, we investigated how non-invasive brain stimulation (10Hz-tACS) alters ANS balance measured with RSA and how these metrics correlated with the level of pain and other self-report characteristics. Contrary to our hypothesis, we found that there was no effect of 10Hz-tACS on RSA. Previous studies using transcranial direct current stimulation increased RSA in healthy participants (57,58). However, we found a significant increase in SDNN for 10 Hz-tACS relative to sham. Our exploratory analyses to find a relationship between baseline RSA and pain severity did not show any significant correlations in agreement with previous findings that included no intervention (59).

While both sequences had a similar tACS effect on RSA (Figure 2A, left panel), we found a trending effect of greater RSA in the tACS condition compared to sham before the intervention at both sessions (Figure 4, F_1,36_=3.105, p=0.08). This finding may have limited the potential influence that 10Hz-tACS had on RSA as there was less potential for an intervention to increase RSA due to ceiling effects (60). Previous evidence suggests that there is an optimal range for RSA based on breathing rate and vagal input (19). Furthermore, most HRV intervention studies including biofeedback training with a slow controlled breathing rate or meditation (61) and HRV changes are measured throughout a longer duration with daily sessions for six weeks (22,24,62). RSA adapts quickly due to both internal and external perturbations (20), therefore a treatment to create a lasting effect needs to be consistent, such as structured resonance breathing training daily (61).

**Figure 4.**
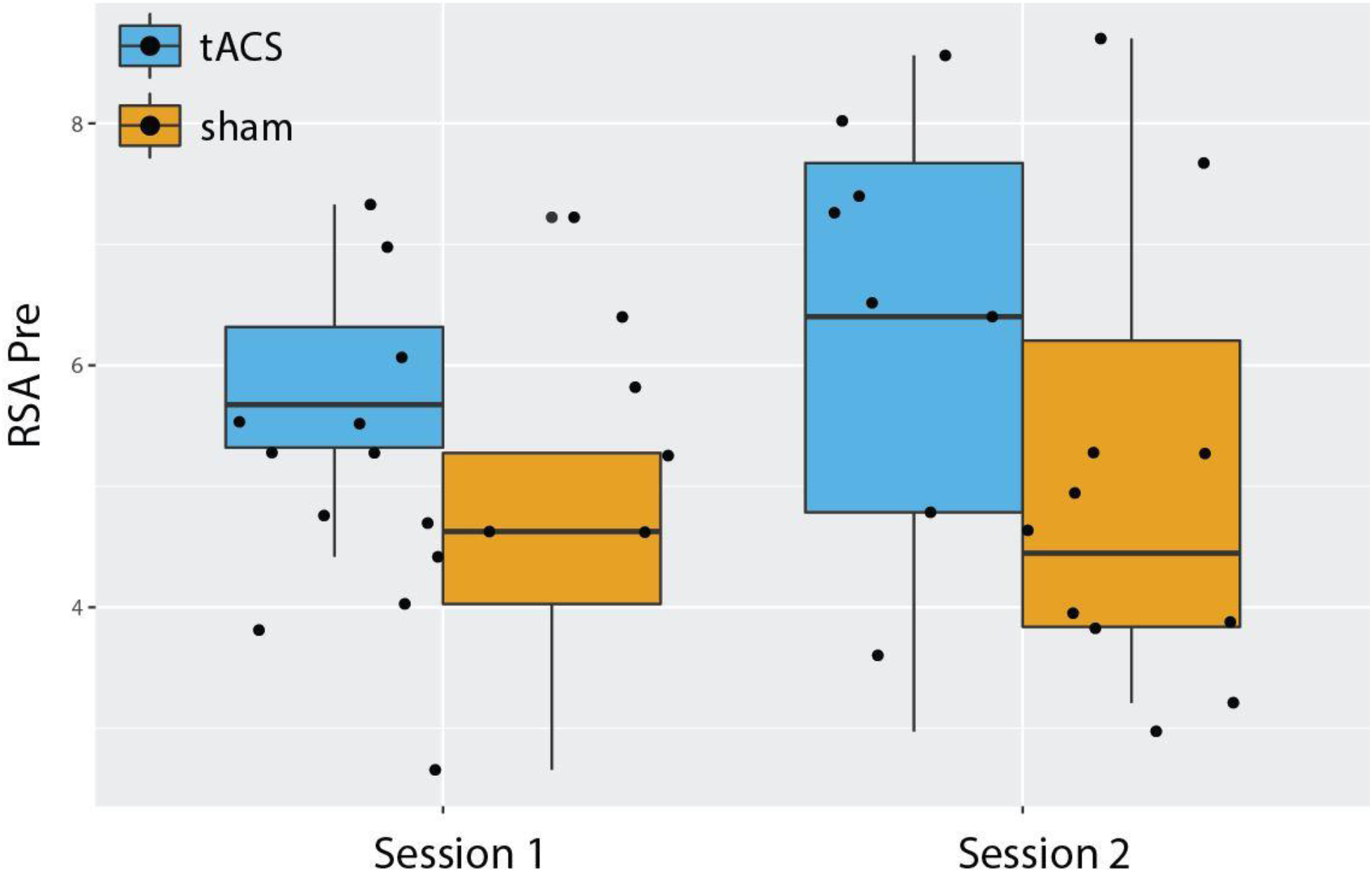
Baseline RSA values before stimulation (tACS and sham) in both sessions.

Within our time-domain analysis, 10Hz- tACS had a significant effect on SDNN. The time domain analysis reports the activity of the cardiac system (63), which may in turn broadly reflect ANS balance (12). SDNN is a commonly used parameter for the measurement of total HRV and represents the overall variability of both sympathetic and parasympathetic inputs to the heart (64). Many studies within chronic pain have found decreased SDNN within clinical populations compared to healthy controls (12) and HRV suppression has been correlated with pain severity or disability perception (59,65,66). One study (67) measured SDNN in patients with spinal cord injury (SCI) with and without neuropathic pain and found significant lower resting SDNN in SCI patients with pain compared to SCI without pain and healthy controls. Since SDNN includes input from both the parasympathetic and sympathetic input, few conclusions on the increase of specific ANS branches can be drawn (12), but SDNN has been hypothesized to provide objective quantification of analgesic response to pain treatment (67). Therefore, increasing overall HRV (SDNN) may be beneficial in patients with CLBP, and HRV intervention studies have shown increases in total HRV (24). Our findings thus suggest that SDNN may be a better target than RSA, at least for a single session of 10Hz-tACS.

Previous studies, which investigated non-invasive brain stimulation interventions in chronic pain, have shown promising, but varied results (57,58,68–70). Most of these studies have focused on transcranial direct current stimulation and transcranial magnet stimulation. Given our findings of successful target engagement of alpha oscillations that correlated with clinical pain improvement as reported in our previous paper (28), tACS has the potential to provide a safe, scientifically-supported, low-cost treatment option. However, more research utilizing tACS is needed to replicate our results and further dissect the underlying mechanism(s). Our detailed characterization (Tables 1 and 2) of the patient population provided here can be used to inform the planning of future tACS studies including the use of power calculations to inform sample size and measures collected.

As with any scientific study, the work presented here has limitations. First, this was a pilot study and thus not designed to establish statistical significance for small effect sizes. Our within-subject design increased our statistical power and is a strength considering large between-subject variation in many HRV components (71). Nevertheless, all statistical results should be interpreted cautiously given the small sample size. Second, we did not collect medication and lifestyle information unrelated to pain, such as antihistamines or caffeine use, both of which have been shown to influence HRV, albeit the within-subjects design should reduce impact of external factors (71). Third, we only analyzed two minutes of ECG data for HRV analysis following other studies in the field (72,73). However, the current gold standard for HRV recordings is at least five minutes (74) and our study may have benefited from longer recordings. Fourth, our crossover study design only allowed for one session of active stimulation. TACS may have an additive effect on modulating oscillations if a design with multiple sessions were used (75,76).

Future studies involving multiple stimulation sessions are the next steps since recurring stimulation sessions are likely needed to produce perceptible and lasting clinical effects due to presumed underlying mechanisms that appear to be related synaptic plasticity (77). Several studies have investigated non-invasive brain stimulation techniques in patients with chronic pain, but treatment effects vary across the studies, and typically only clinical outcomes are reported. We aimed to identify objective biological targets using EEG and ECG to better understand the action of 10Hz-tACS and the role of the ANS in chronic pain. Our results presented here along with those in our previous paper (28) show that brain network dynamics and self-reported pain seem to be more sensitive ways than HRV metrics to measure effects of brain stimulation for individuals with CLBP.

## Acknowledgements

This work was supported by the National Institute of Mental Health of the National Institutes of Health under Award Numbers R01MH111889 and R01MH101547, the National Center for Advancing Translational Sciences (NCATS), National Institutes of Health, through Grant Award Number UL1TR002489. The content is solely the responsibility of the authors and does not necessarily represent the official views of the NIH. We gratefully acknowledge the help and support from the Carolina Center for Neurostimulation.

## Author contributions

All authors have read and approved to submit the paper. F.F., J.H.P., and K.L.M designed the study. J.H.P. and M.L.A. performed the study. J.H.P., M.D, and S.A. analyzed the data. J.H.P and F.F. wrote the manuscript and all authors contributed to editing the manuscript.

